# Novel *Streptococcus mitis* subsp. *mitis* Picklauza buccal isolate from individual with idiopathic pulmonary fibrosis containing functional mobile *qacG* Gene

**DOI:** 10.64898/2026.10.15.759443

**Authors:** Sydnie N. Klauza, Sophie M. P. Pickelner, Kelyah Spurgeon, Souzane Ntamubano, Nicole R. Falkowski, Nisha Mohan, Jay Lipinski, Justin M. Oldham, David N. O’Dwyer, Ariangela J. Kozik

## Abstract

Idiopathic pulmonary fibrosis (IPF) is a chronic, progressive lung disease characterized by immune dysregulation, fibrosis, and eventual respiratory failure. While the etiology of IPF remains unclear, accumulating evidence implicates the oral and airway microbiome in disease progression, with elevated Streptococcal burden linked to increased mortality. Whether specific taxa actively contribute to IPF pathophysiology or arise secondarily from the damaged lung environment remains unresolved. Here, we characterize “Picklauza,” a streptococcal strain isolated from the buccal mucosa of an individual with IPF. Although its average nucleotide identity to a *Streptococcus mitis* reference genome was 93.855%, analysis with Streptoquod supported classification as *Streptococcus mitis* subsp. *mitis*. Genomic analysis identified a 109-amino-acid *qacG* homolog predicted to encode a small multidrug resistance (SMR)-family efflux transporter. The locus was located near genes associated with mobile genetic elements, raising the possibility of horizontal acquisition. The predicted QacG protein retained residues conserved in the extensively characterized SMR transporter EmrE, and RT-qPCR detected *qacG* transcripts under standard culture conditions. Nevertheless, Picklauza did not display increased tolerance to benzalkonium chloride, benzethonium chloride, or ethidium bromide compared with *S. mitis* NCTC 12261. Picklauza also differed from the reference strain in growth kinetics, carbon utilization, and antimicrobial susceptibility. PCR screening detected *qacG* homologs in three additional oral isolates from patients with IPF. Together, these findings identify a transcriptionally active, putatively mobile *qacG* homolog in oral streptococci while revealing a disconnect between gene carriage and QAC susceptibility under the conditions tested. This work highlights the need to validate predicted resistance determinants phenotypically and to investigate the mobility and conditional activity of SMR-family genes across genus boundaries in host-associated microbial communities.

**Importance:** Genomic surveillance frequently infers antimicrobial resistance from the presence of annotated resistance genes, but gene carriage does not always predict phenotype. We characterized *Streptococcus mitis* subsp. *mitis* strain Picklauza, isolated from the buccal mucosa of an individual with IPF, and identified a *qacG* homolog predicted to encode an SMR-family efflux transporter. Although the locus was transcriptionally active and retained conserved residues associated with SMR transport, Picklauza remained susceptible to the tested QACs and ethidium bromide. Related *qacG* sequences were also detected in additional IPF associated oral isolates. These results illustrate the importance of pairing genomic surveillance with functional testing and provide a foundation for investigating the mobility, regulation, and substrate specificity of SMR-family genes in oral streptococci.

## INTRODUCTION

### Idiopathic pulmonary fibrosis and host-associated microbiota

Idiopathic pulmonary fibrosis (IPF) is a chronic, progressive lung disease characterized by fibrosis, declining pulmonary function, and high mortality. IPF affects an estimated 3 million people worldwide, although reported prevalence varies substantially across populations and diagnostic criteria. Median survival following diagnosis is approximately 3 to 5 years. Available antifibrotic therapies can slow disease progression, but they do not reverse established fibrosis, and currently lung transplantation remains the only definitive treatment for eligible patients^1–3^.

Although the cause of IPF remains unknown, growing evidence links respiratory microbial communities to disease severity and progression. Increased airway bacterial burden and altered community composition have been associated with a rapid functional decline, acute exacerbations, and mortality^4^. Streptococci are among the taxa frequently detected in the upper and lower respiratory tracts, and enrichment of particular streptococcal lineages has been associated with adverse clinical outcomes^4,5^. Whether these organisms contribute directly to disease progression or their populations expand in response to an altered airway environment remains unresolved.

The oral cavity is a potential source of microbes entering the lower respiratory tract through microaspiration. Characterizing oral isolates from patients with IPF may therefore help define microbial traits that facilitate persistence across connected oral and respiratory niches. However, the oral microbiota of patients with IPF and the physiology of individual patient-derived isolates remain comparatively understudied.

Antibiotics, such as Co-trimoxazole, a combination of trimethoprim and sulfamethoxazole^6^, are often used in an IPF treatment course, especially when bacterial infection or acute exacerbation is suspected. Azithromycin, a macrolide antibiotic, may be also used long term at low doses for anti-inflammatory properties. Such exposure can alter microbial community structure and select for antimicrobial resistance, but its effects on oral and respiratory commensals remain poorly defined. Studying cultured isolates alongside community level data can help connect genomic predictions to physiological phenotypes and clarify how treatment-associated pressures shape host-associated bacteria.

### Quaternary Ammonium compounds and efflux-mediated tolerance

Patients receiving frequent healthcare may encounter both antibiotics and disinfectants. Quaternary ammonium compounds (QACs), including benzalkonium chloride (BAC) and benzethonium chloride (BZT), are cationic surfactants used in clinical, household, and personal-care products. Their use increased during the COVID-19 pandemic, raising questions about how repeated or subinhibitory exposure affects host-associated and environmental microbial communities ^7–10^.

QAC tolerance can be mediated by multidrug efflux systems ^11^. Small multidrug resistance (SMR)- family transporters are compact membrane proteins, typically approximately 100–120 amino acids long, that contain four transmembrane helices and function as oligomeric proton-coupled transporters ^12–15^. The *Escherichia coli* transporter EmrE is a well-characterized member of this family. Conserved residues, including Glu-14, contribute to proton coupling and substrate transport, whereas other residues help determine folding, oligomerization, and substrate recognition ^15,16^. These efflux pumps export toxic molecules, typically conferring increased tolerance to an assortment of substances containing these compounds ^12^ such as disinfecting agents, intercalating dyes, such as ethidium bromide and on occasion antibiotics such as certain beta-lactams, aminoglycosides, and fluoroquinolones ^7,13^. QACs are effective against Gram-positive and Gram-negative bacteria and are frequently used in biocidal and personal care products ^12^. QAC concentrations below minimal inhibitory concentrations (MIC) can have different effects on bacterial cells, sometimes increasing DNA mutation rates and the transfer of resistance genes ^17^.

The nomenclature of QAC-associated efflux genes is complex. Genes commonly designated *qacC*/*qacG* encode SMR-family proteins, whereas *qacA* and *qacB* encode the much larger major facilitator superfamily (MFS) transporters. These similarly named genes therefore belong to distinct transporter families and should not be treated as functionally interchangeable ^11,15,18,19^. SMR-family proteins were originally described in *Staphylococcus aureus*^18^, but *emrE* in *Escherichia coli* has been studied extensively and now serves as the reference point for these genes ^15,19^. SMR genes are frequently associated with plasmids and other mobile genetic elements, which may facilitate their dissemination ^14,19^.

### Horizontal gene transfer and mobile resistance determinants

Horizontal gene transfer allows bacteria to acquire genetic material through transformation, transduction, or conjugation and is a major driver of microbial adaptation ^20–26^. Resistance determinants can move not only among closely related strains but also across genera, particularly when carried by plasmids, integrons, transposons, or integrative and conjugative elements (ICEs). For example, QAC-associated genes historically linked to staphylococci have been reported in other Gram-positive genera, but rarely in Streptococci ^27–29^.

Mobile genetic elements can accumulate resistance and stress-response genes and may facilitate their persistence under antimicrobial selection. However, the presence of a resistance-associated gene does not guarantee a measurable resistance phenotype. Functional expression may depend on promoter context, transcript abundance, translation, membrane insertion, oligomerization, energetic coupling, and substrate specificity. Consequently, genomic annotation should be paired with phenotypic testing.

### Knowledge gap and study aims

QAC-associated SMR genes are well described in staphylococci but remain poorly characterized in oral streptococci. In a companion Microbiology Resource Announcement, we reported the genome of Picklauza, a streptococcal strain isolated from a buccal swab collected from a patient with IPF. Preliminary genomic analysis identified a *qacG* homolog within a region containing genes associated with mobile genetic elements.

Here, our objective was to characterize Picklauza in comparison with the type strain *S. mitis* NCTC 12261 (ATCC 49456). We assessed colony and cellular morphology, growth kinetics, carbon utilization, antimicrobial susceptibility, QAC and ethidium bromide susceptibility, and basal *qacG* transcription. We also examined the predicted *qacG* sequence and structure, analyzed the surrounding genomic region, and screened additional oral isolates from IPF patients for related *qacG* sequences. We hypothesized that carriage and expression of *qacG* would be associated with increased tolerance to QACs. Instead, Picklauza remained sensitive under the tested conditions, revealing a genotype–phenotype discrepancy that emphasizes the need for functional validation of predicted resistance genes.

## Materials and Methods

### Strain isolation and identification

Buccal swabs were collected from patients with IPF at the University of Michigan Health System (IRB HUM00105694). Strain isolation was performed as described in the companion Microbial Resource Announcement. Gram staining was performed to identify cellular morphology and Gram reaction.

Because Sanger sequencing and targeted colony PCR did not provide a definitive taxonomic assignment, the isolate was submitted to Plasmidsaurus for whole-genome sequencing. A 5-mL culture was grown overnight in brain heart infusion (BHI) broth at 37°C under aerobic conditions. Cells were harvested by centrifugation, washed twice with 1 mL of 1X phosphate-buffered saline, and resuspended in 500 µL of DNA/RNA Shield (Zymo Research). Sequencing, genome assembly, taxonomic analysis, and phylogenetic methods are described in the companion report.

### Culture conditions

*Streptococcus mitis* NCTC 12261 was purchased from ATCC (ATCC 49456) as a lyophilized culture. Strain Picklauza and NCTC 12261 were grown and maintained aerobically using Brain Heart Infusion (BHI) agar supplemented with 5% sheep’s blood and BHI broth at 37°C. Hemolysis was observed after growing Picklauza on BHI agar with 5% sheep’s blood.

### Capsule staining

Capsule staining was performed using Anthony’s staining method from ASM ^30^. Milk broth was prepared as described using Lab Scientific Non-Fat dry milk (Fisher Scientific NC9121673) ^30^. 5 mL milk broth cultures of Picklauza and NCTC 12261 were grown at 37°C for 15 hours. A 10 µL smear of each culture was prepared on a blank glass slide. The smears were stained for 2 minutes with 1% crystal violet and washed gently with 20% copper sulfate. Images were taken at 1000X magnification with a Nikon E800. We prepared and viewed 6 independent cultures on 10 slides at 12, 16 and 18 hours of growth in milk broth.

### Growth kinetics and CFU

Overnight cultures were diluted in fresh BHI broth to an initial OD_600_ of 0.05. Aliquots of 200 µL were transferred to a flat-bottom 96-well plate, with six biological replicates per strain. Plates were incubated aerobically at 37°C in an Alto plate reader (Cerillo), and optical density was measured every 10 min for 24 h. Blank-corrected growth curves were used to calculate doubling times during exponential growth using Labrador software (Cerillo).

Samples for CFU plating were taken at the start and end time points from all replicates of Picklauza and NCTC 12261. Each sample was serially diluted tenfold in water to a dilution factor of 10^-11^. Three technical replicates of each dilution were plated in 10 µL spots on BHI plates with 5% sheep’s blood and grown aerobically at 37°C for 24 hours. Colonies were counted in the first easily countable spot and averaged between the technical replicates. Resulting colony numbers were plugged into the following equation and then log transformed 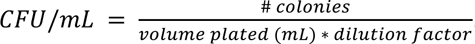 Welch’s *t*-test statistical analyses were used to determine statistical significance (p<0.05).

### Assaying Antibiotic Susceptibility and Carbon Utilization

Antibiotic susceptibility profiles were generated using the BIOLOG^©^ Phenotype MicroArray (PM) bacterial chemical sensitivity panels, PM11C and PM12B. These plates contain lyophilized antibiotics and chemicals, present in four concentrations each, to allow for the dose-response assessment of 48 total substrates. 5 mL BHI broth cultures of each strain (Picklauza and NCTC 12261) were grown overnight at 37°C under aerobic conditions. New BHI cultures with a starting OD of 0.05 were prepared from the overnight cultures and added to PM11C and PM12B at a volume of 150 µL per well. Untreated positive control conditions of Picklauza and NCTC 12261 were added to empty BIOLOG^©^ 96-well plates. Growth was measured in an ODIN^©^ plate reader at OD740, with measurements taken every 20 minutes. The area under the curve (AUC) value for each antibiotic class across the four concentrations was averaged to generate antibiotic sensitivity groupings.

Carbon utilization was assessed using the BIOLOG^©^ Phenotype MicroArray (PM) panels PM1 and PM2. Combined, the two panels contain 190 unique lyophilized carbon sources and an empty negative control well (A1). Cultures of Picklauza and NCTC 12261 were grown overnight at 37°C in BHI broth. Inoculating cultures with a starting OD of 0.05 were made in a modified *Streptococcus sp.* Inoculating Fluid from Biolog (In 100 mL: 2 mM MgCl2, 1 mM CaCl2, 0.6 yeast extract, and 0.12 mL Dye Mix G (Biolog)). Cultures were added to PM1 and PM2 at a volume of 150 µL per well and growth was measured in an ODIN^©^ plate reader. OD was measured at 590 and 740 nm, but analysis was restricted to 740 nm readings to prevent interference from substrate absorbance. To generate ΔAUC values, the AUC from the negative control well was subtracted from AUC values for each substrate.

### Quaternary ammonium compound and ethidium bromide susceptibility testing

Susceptibility to benzalkonium chloride (BAC; Sigma-Aldrich, 12060), benzethonium chloride (BZT; Sigma-Aldrich, B8879), and ethidium bromide (EtBr; Sigma-Aldrich, 1003903523) was assessed using concentration-dependent survival assays. BAC and BZT were dissolved in sterile water to 10% (w/v) and passed through 0.2-µm syringe filters. EtBr was purchased as a 1% (w/v) solution.

Picklauza and *Streptococcus mitis* NCTC 12261 were grown overnight in 5 mL brain heart infusion (BHI) broth at 37°C under aerobic conditions. Overnight cultures were diluted into fresh BHI broth to a starting optical density of 0.05. For BAC assays, cultures were exposed to final BAC concentrations of 0.0001%, 0.005%, 0.01%, 0.05%, 0.1%, 0.5%, or 1% (w/v). Controls included uninoculated BHI, an untreated culture of each strain in BHI, and uninoculated BHI containing 1% BAC. Assays were conducted in polypropylene tubes to minimize adsorption of the cationic compounds to negatively charged polystyrene surfaces ^31^. Cultures were incubated aerobically for 6 h at 37°C.

Samples were collected immediately after compound addition (0 h) and after 6 h of exposure. Viable-cell concentrations were determined by serial dilution and plating as described above. Colonies were counted after 24 h of incubation, and plates were incubated for an additional 48 h to assess hemolytic activity. CFU/mL values were log_10_-transformed before analysis.

BZT susceptibility testing was performed using the same procedure and exposure period as the BAC assay, with final BZT concentrations of 0.0001%, 0.005%, 0.01%, 0.05%, 0.1%, 0.5%, or 1% (w/v). EtBr susceptibility testing was performed with final concentrations of 0.0001%, 0.005%, 0.01%, 0.05%, and 0.1% (w/v). EtBr-exposed cultures were incubated aerobically for 24 h at 37°C before viable-cell concentrations were determined by serial dilution and plating.

Concentration-dependent survival was evaluated by comparing viable-cell concentrations across treatment conditions. Statistical analyses were performed in GraphPad Prism using one-way ANOVA and Welch’s *t*-test, followed by Tukey’s multiple-comparison test. Statistical significance was defined as *P* < 0.05.

### DNA and RNA extraction

Genomic DNA was extracted using the DNeasy Blood and Tissue Kit (Qiagen). RNA was extracted using the Qiagen RNAprotect Bacteria Reagent Handbook on overnight cultures of NCTC 12261 and Picklauza grown in BHI broth. RNA was treated with DNase and assessed for concentration and purity using NanoDrop Lite (Thermo Scientific). cDNA was made following the RT2 Profiler PCR Array Handbook.

### Real Time Quantitative PCR (RT-qPCR)

Primers were ordered from IDT as described in Table 1. To optimize primers, gDNA was isolated from NCTC 12261 and Picklauza at a concentration of roughly 200 ng/uL. Primers were tested at a range of temperatures (52°C, 57°C, and 62°C) and concentrations (200 nm and 400 nm). 62°C and 400 nm were selected as optimal conditions for qPCR of cDNA.

**Table 1.**
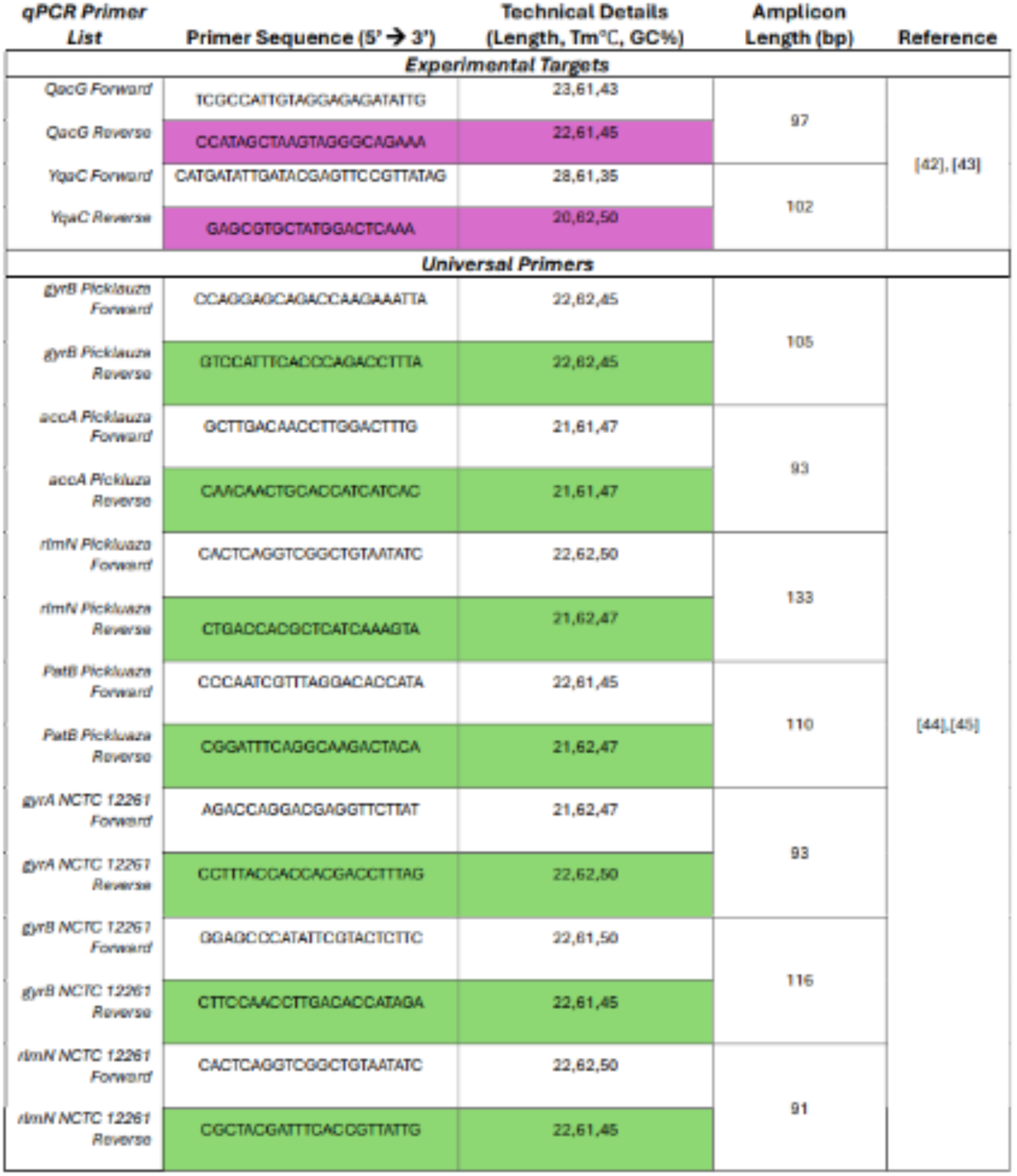
Primer sequences and specifications for Picklauza and NCTC 12261. Experimental primers for *qac* genes are highlighted in purple, while housekeeping genes for both are highlighted in green.

Quantitative PCR assays were conducted in a 20 μL final reaction volume per well using PowerTrack SYBR Green Master Mix (Applied Biosystems). Each reaction consisted of 10 μL of 2X master mix, 4 μL of primer mix (forward and reverse; 400 nM final concentration of each primer), 0.5 μL of PowerTrack Yellow Sample Buffer, 0.5 μL of nuclease-free H₂O, and 1 μL of cDNA template (200 to 400 ng/μL, depending on the replicate). Amplification and detection were performed on a QuantStudio 3 Real-Time PCR System (Applied Biosystems). The thermal profile consisted of 40 cycles of denaturation at 95°C for 15 seconds, annealing at 62°C for 30 seconds, and extension at 72°C for 1 minute. To confirm amplification specificity, a melt curve analysis was performed immediately following the amplification cycles by heating to 95°C for 15 seconds, cooling to 60°C for 1 minute, and then increasing the temperature to 95°C for 15 seconds. Amplification efficiencies were calculated from standard curves. Transcript abundance was normalized to gyrB, gryA, accA, rlmN, and patB using the ΔCt method. ΔCt was calculated by taking the average of each technical replicate. Genomic mean of all housekeeping genes was taken and subtracted from the *qac* genes of interest. 3 biological replicates with 3 technical replicates each were included.

## RESULTS

### Picklauza displays alpha hemolysis, an extracellular capsule, and aggregation in liquid culture

Primary isolation from a buccal swab collected from a patient with idiopathic pulmonary fibrosis (IPF) yielded pure colonies of strain Picklauza following serial passage. Colonies appeared convex, white to gray in color, with irregular, lobate colony margins. Picklauza colonies also displayed alpha hemolysis when grown on 5% sheep blood agar (Figure 1A). Microscopic evaluation via Gram staining confirmed the expected cellular morphology of Gram-positive streptococci (Figure 1B) and capsule staining using Anthony’s capsular staining method confirmed an extracellular polysaccharide capsule surrounding the cell wall of Picklauza (Figure 1C). In liquid media, stationary overnight cultures of strain Picklauza formed visible aggregates, whereas NCTC 12261 remained more uniformly suspended (Figure 1D).

**Figure 1.**
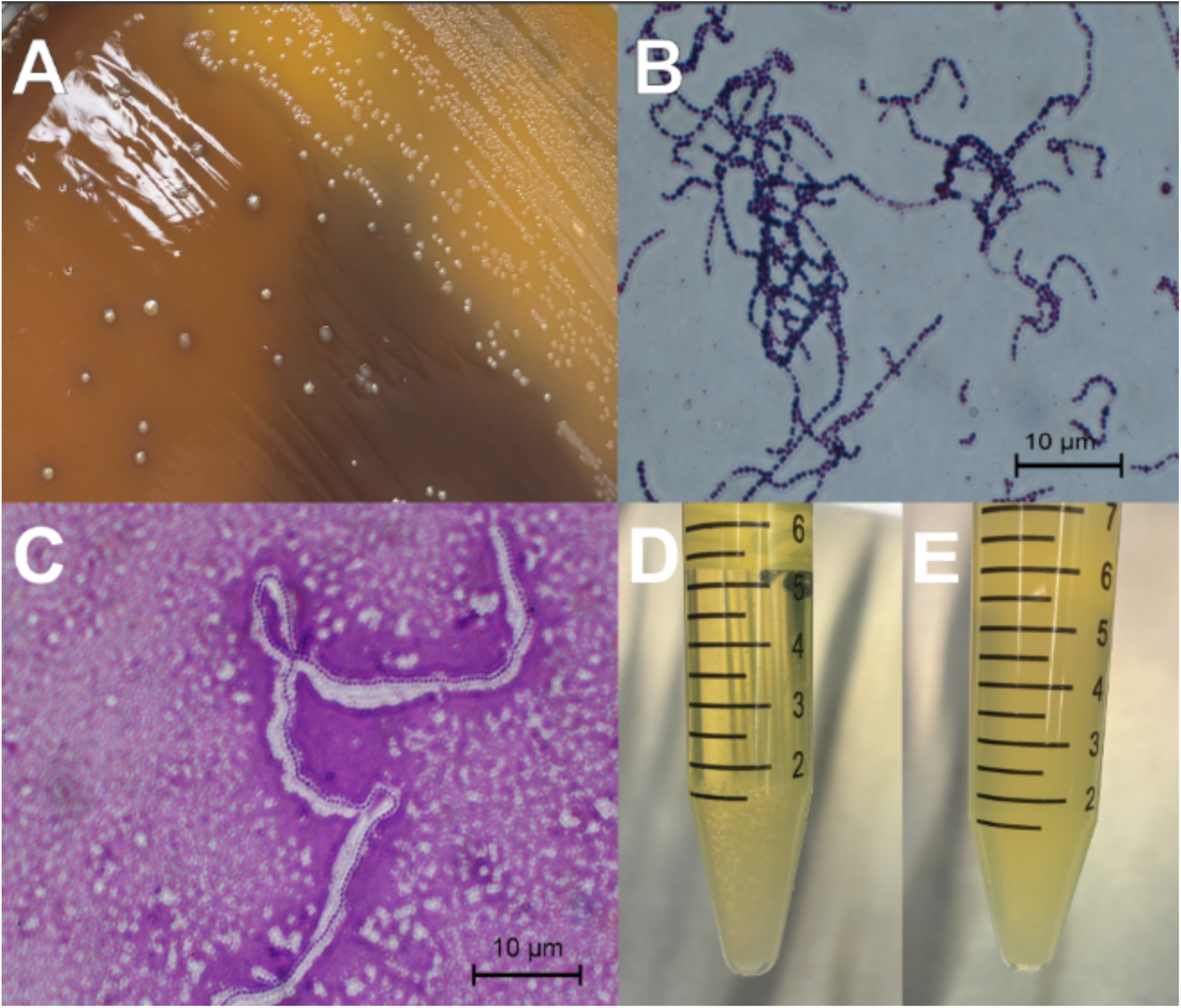
Phenotypic characterization of Picklauza. (A) Colony morphology and hemolytic activity of Picklauza cultured on BHI blood agar. (B) Light microscopy image of gram stained Picklauza cells, gram positive 10 µm. (C) Light microscopy image of capsule stained Picklauza displaying large clear halo against stained crystal violet background (10 µm). (D) Picklauza overnight culture grown in BHI broth. (E) NCTC 12261 overnight culture grown in BHI broth.

### Strain Picklauza exhibits a reduced growth rate while maintaining equivalent final biomass and cell viability to the type strain

To evaluate whether the observed phenotypic differences in liquid media impacted overall replication kinetics, we monitored growth curves and viable cell yields over 24 hours alongside the NCTC 12261 reference type strain. Both strains attained a similar maximum optical density (OD600) over the 24-hour time course (Figure 2A). However, strain Picklauza exhibited a distinct shift in growth kinetics, replicating more slowly with a calculated doubling time of 1.51 hours compared to the type strain (0.8 hours). Despite this slightly altered growth kinetic profile, final viable cell counts based on colony forming units (CFU/mL) following 24 hours of incubation revealed no statistically significant differences between the two strains (Figure 2B).

**Figure 2.**
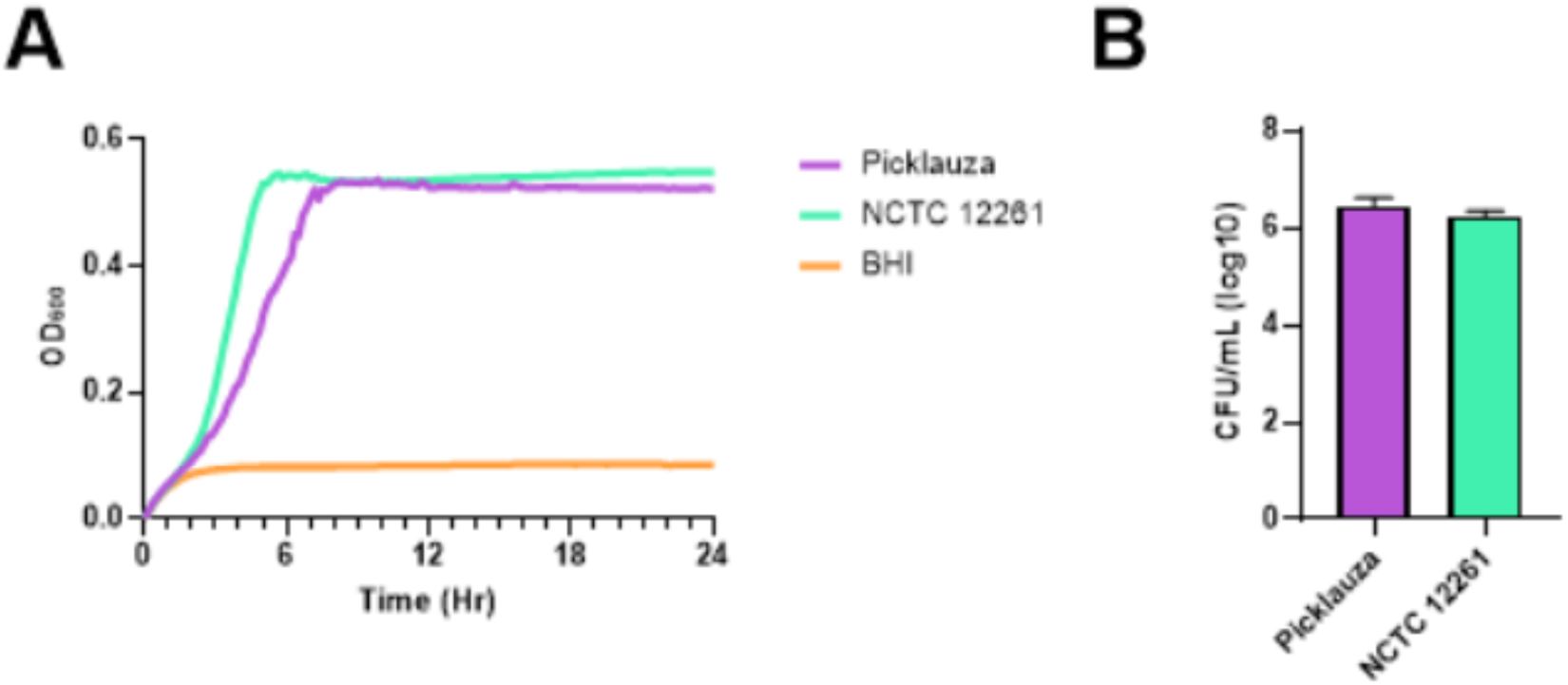
Growth dynamics and viability of Picklauza and NCTC 12261. (A) Aerobic growth curves of NCTC 12261 and Picklauza in BHI broth over 24 h (n = 6). (B) Log transformed colony forming unit (CFU/mL) counts from endpoint of aerobic growth curves (24 h). Data represented as the mean of three technical replicates from six biological replicates.

### Picklauza and NCTC 12261 display distinct responses across carbon-source conditions

To assess whether morphological and kinetic differences reflected broader metabolic shifts, we conducted high-throughput metabolic profiling of >180 carbon sources that revealed substantial divergence between strain Picklauza and NCTC 12261 (Figure 3 and supplemental figure 1). Both strains demonstrated robust metabolic activity on core carbohydrate substrates; however, strain Picklauza displayed a distinct substrate utilization profile compared to NCTC 12261. Specifically, within the simple carbohydrate and carboxylic acids groups (Figure 3), Picklauza and *NCTC 12261* have unique growth patterns. Both strains responded strongly to several common carbohydrate substrates, but they differed in wells containing maltotriose, D-fructose, D-mannose, glycogen, D-cellobiose, D-fucose, and propionic acid. These findings indicate strain-level differences in metabolic activity or growth under the tested conditions. Additional targeted assays will be required to determine whether the observed differences reflect substrate catabolism, differential tolerance, or indirect effects of the assay medium.

**Figure 3.**
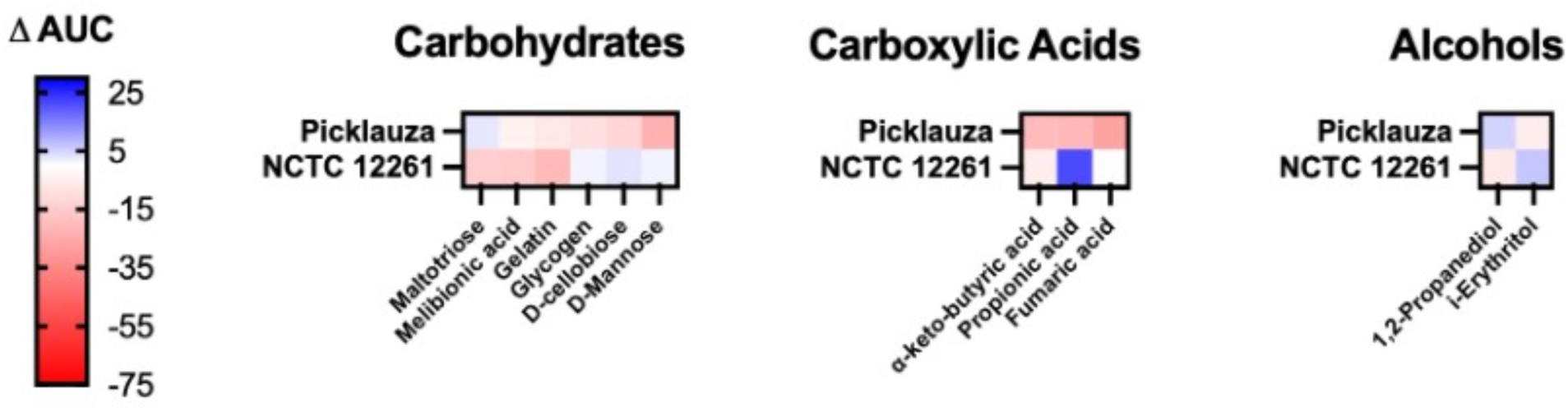
Heatmap containing most significant differentially utilized carbon sources between NCTC 12261 and Picklauza. Picklauza and NCTC 12261 demonstrate differential utilization of several carbohydrates, carboxylic acids, and alcohols. Significantly different utilization determined as differences between ΔAUC values of >10 or <-10. ΔAUC calculated by subtracting the negative control from the average AUC value for each carbon source (n=3). Cells inoculated in *Streptococcus* mixture IF and adjusted to an OD600 of 0.05 prior to measuring growth for 24 hours at OD740. Carbon sources are hierarchically ordered from highest relative ΔAUC (blue) to lowest relative ΔAUC (red) based on the utilization profile of Picklauza.

### High-throughput antimicrobial profiling reveals broad antibiotic susceptibility alongside selective resistance in strain Picklauza

To establish the drug resistance landscape of Picklauza, we evaluated its sensitivity against a high-throughput panel of standard antimicrobial agents alongside NCTC 12261 (Figure 4 and supplemental table 2). Strain Picklauza remained susceptible to the core first-line of therapeutic antibiotic beta-lactams, including penicillin and cephalosporin. However, distinct resistance profiles emerged between the two strains, with strain Picklauza displaying elevated resistance toward rifamycin, macrolides, and amphenicols. Uniquely, our preliminary screening highlighted an unexpected tolerance toward non-antibiotic antiseptic compounds setting the stage for genomic investigation into specific resistance determinants.

**Figure 4.**
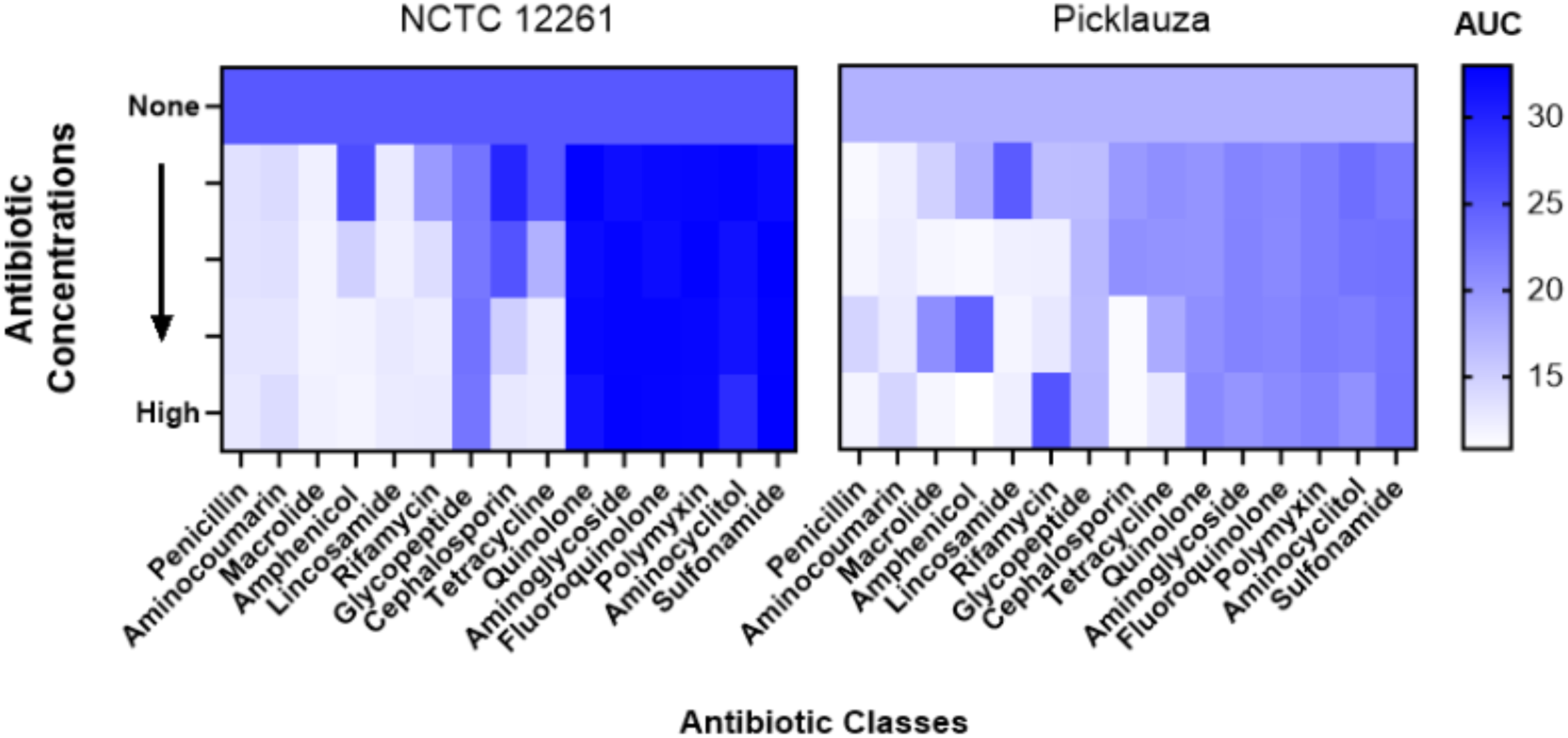
Antimicrobial susceptibility profiles of Picklauza and NCTC 12261. Antibiotic growth profiles determined by AUC measurements across four different concentrations (n = 3). BHI cultures were inoculated to an initial optical density (OD600) of 0.05, and growth was measured for 24 hours at OD740. High relative AUC values (dark blue) indicate robust ability for growth within specified antibiotic class, while decreasing relative values (blue to white) denote growth inhibition relative to the untreated control group for both strains.

### Picklauza encodes a *qacG* homolog with residues conserved among SMR-family transporters

Targeted genomic analysis of the strain Picklauza assembly identified a small multidrug resistance (SMR) family gene, designated *qacG*, predicted to encode a quaternary ammonium compound efflux protein. To evaluate whether this locus represents a functional transporter, we aligned the predicted *qacG* amino acid sequence from strain Picklauza with the canonical SMR efflux pump EmrE from *Escherichia coli* (Figure 5). The alignment revealed identical conserved charged residues Glu-14, Lys-22, and His-110. This also revealed conserved substrate binding residues Tyr-40 and Trp-63.

**Figure 5.**
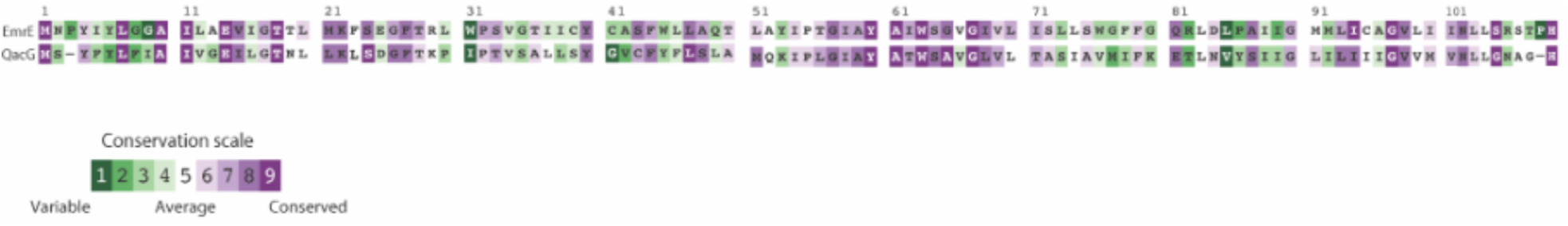
Alignment of conserved residues in EmrE and Picklauza QacG. Amino acid sequence alignment comparing Picklauza *qacG* protein with the E. coli *EmrE* multidrug transporter. Color coded sequence illustrates position specific conservation on scale from 1 to 9.

The darkest shade of green (1) denotes highly variable amino acid positions, whereas dark purple (9) denotes most structurally and functionally conserved residues within the *qacG* sequence.

To further validate whether the Picklauza *qacG* gene encodes a structurally viable SMR transporter, we generated three-dimensional structural models of *E. coli emrE* and Picklauza *qacG* to illustrate both structural and sequence similarity in the tertiary conformation (Figure 6). To confirm the helical formation similarity between Picklauza *qacG* (pink) and *emrE* (blue), we superimposed the three-dimensional structures (Figure 6).

**Figure 6.**
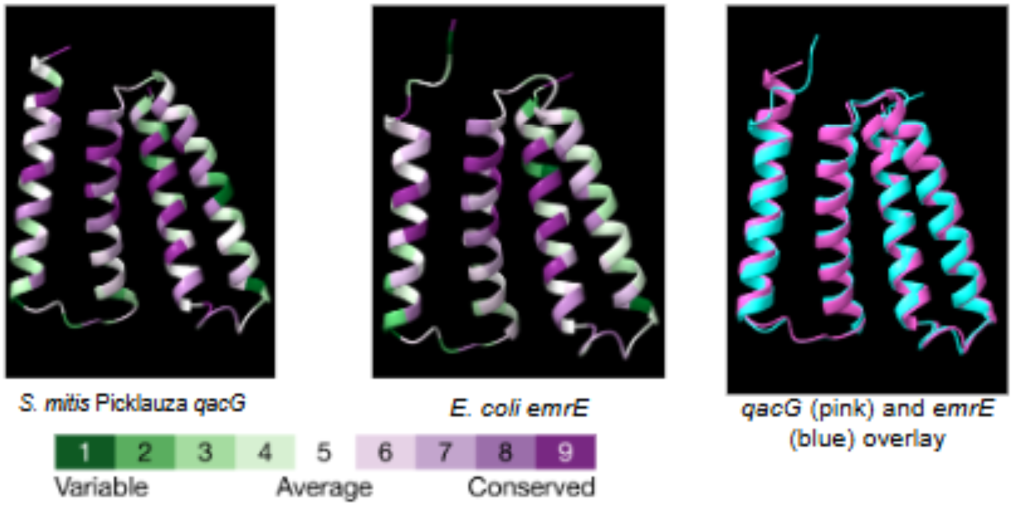
Comparison of 3D conformations of EmrE and Picklauza QacG. Three-dimensional protein structure alignment of Picklauza *qacG* efflux pump against E. coli *EmrE* multidrug transporter in ChimeraX^32^. Color coded sequence from 1 to 9 denotes conservation of structure. Darkest shade of green (1) indicating highly variable surface/ structure of amino acids, while dark purple (9) delineates highly conserved residues critical for functional *qacG* gene.

### The Picklauza *qacG* locus is flanked by mobile genetic element markers, suggesting horizontal acquisition via an integrative and conjugative element

To investigate the potential evolutionary origin and mobility of *qacG* in Picklauza, we analyzed the local genomic architecture surrounding the locus (Figure 7). Synteny mapping revealed that *qacG* resides within a genomic region containing several genes associated with horizontal gene transfer and mobile genetic elements (MGEs). Specifically, an integrative and conjugative element (ICE) integrase (ICEBs1) gene and a putative DNA relaxase (NicK) gene were located in the immediate vicinity of *qacG*. Additional neighboring genes included DNA-binding protein G5P and a chromosome anchoring protein RacA.

**Figure 7.**
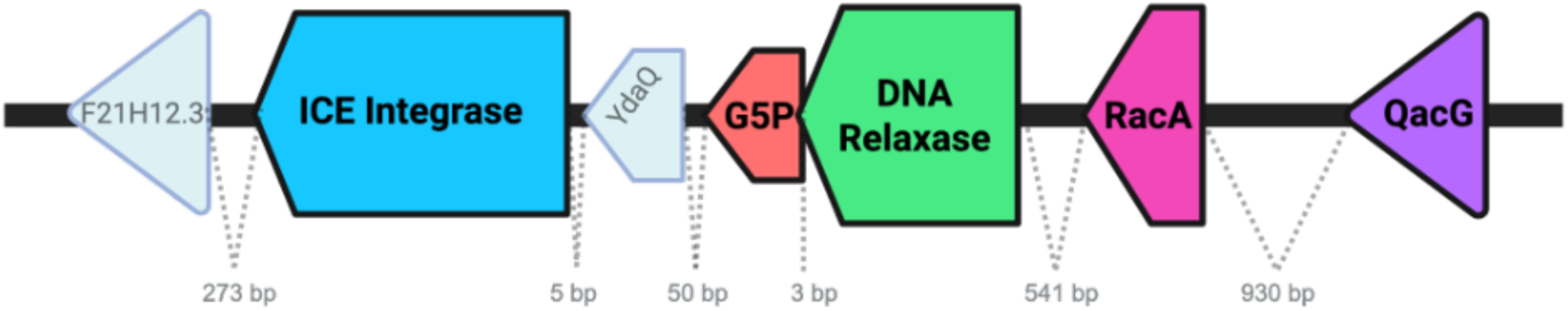
Local genomic map and architectural organization surrounding *qacG* in Picklauza. Linear map illustrates genomic neighborhood of *qacG* locus. Intergenic distances separating adjacent coding regions are indicated in grey text and measured in base pairs (bp). Surrounding downstream and upstream loci consist predominantly of genetic elements often associated with HGT.

### Functional assays reveal retained sensitivity of strain Picklauza to quaternary ammonium compounds and ethidium bromide despite carriage of *qacG*

Despite carrying a *qacG* homolog, Picklauza did not exhibit greater survival than NCTC 12261 following exposure to BAC, BZT, or EtBr (Figure 8). Picklauza showed lower tolerance to EtBr than NCTC 12261 under the tested conditions, with growth seen only at 0.005% and below, whereas NCTC 12261 showed tolerance in up to 0.01% EtBr. The strains exhibited similar responses to BZT, with no growth seen above concentrations of 0.001%. Responses to BAC differed between strains, with no growth of Picklauza observed at 0.005%. The pattern observed for NCTC 12261 requires additional testing with narrower concentration intervals before a definitive endpoint can be assigned. At 0.005% NCTC 12261 exhibited no growth, but grew in concentrations up to 0.05%, and had sustained reduced viability at 0.01% BAC compared to the non-treated control. Thus, carriage of *qacG* was not associated with increased tolerance to the tested compounds under these assay conditions.

**Figure 8.**
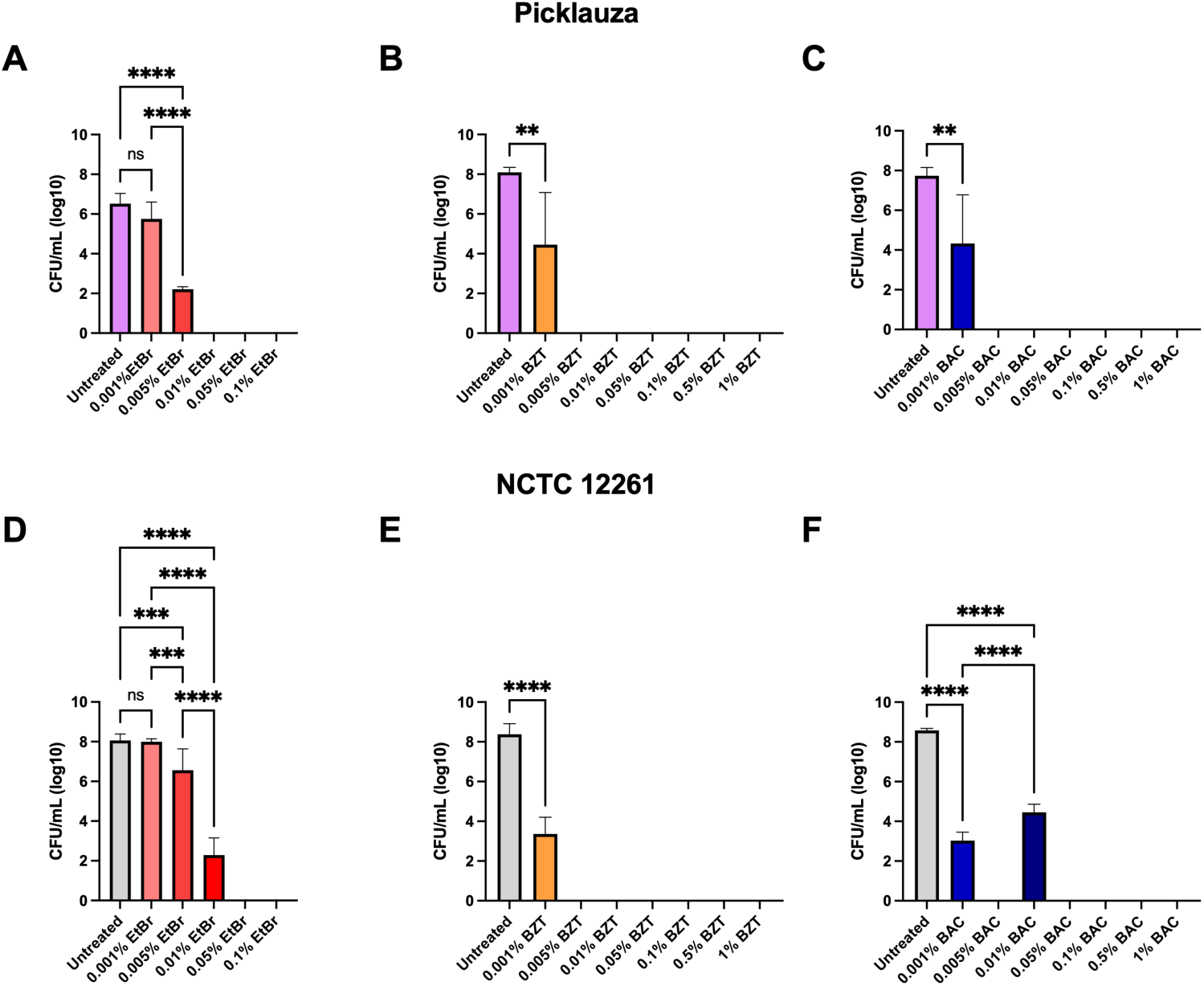
Sensitivity of Picklauza and NCTC 12261 to QACs and EtBr. Graphs display log transformed CFU/mL values for Picklauza (A-C), and NCTC 12261 reference strain (D-F). Strains were treated with 0.001%-0.1% EtBr (A & D), 0.001%-1% BZT (B & E), and 0.001%-1% BAC (C & F). Statistical significance for differences in cell viability was determined using one-way ANOVA and Welch’s *t*-test, followed by Tukey’s multiple-comparison test. Statistical significance was defined as *P* < 0.05.

### Picklauza qacG is transcribed under standard culture conditions

Quantitative RT-PCR using normalized housekeeping controls *gyrA, gyrB, rlmN,* and *patB* for Picklauza demonstrated that both *Yqac* in NCTC 12261 and *qacG* in strain Picklauza are actively transcribed during logarithmic and early stationary phases (Figure 9). Expression levels remained stable across experimental replicates, confirming that the lack of QAC hyper-resistance is not due to transcriptional alterations.

**Figure 9.**
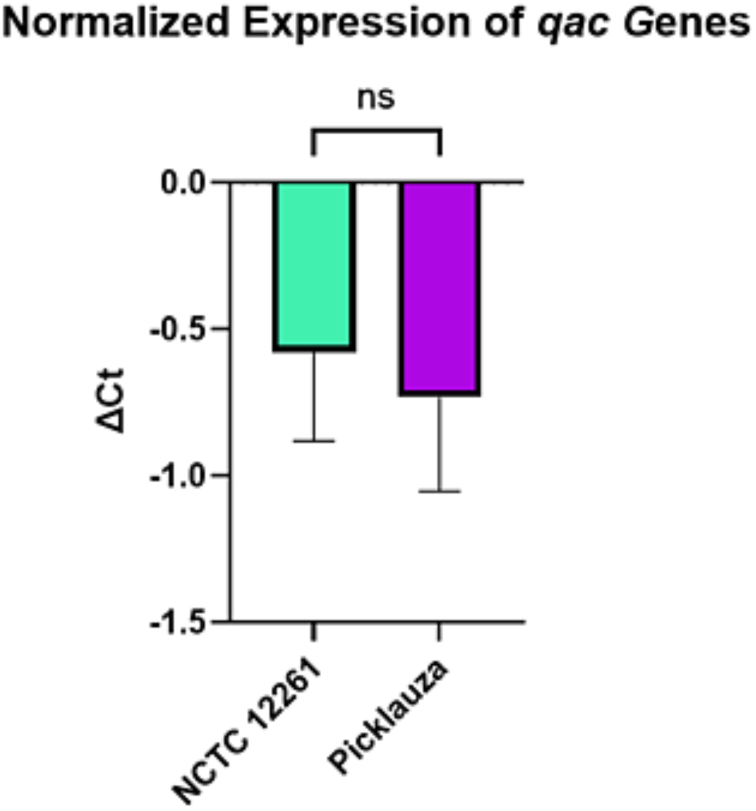
ΔCt levels of *qac* genes in Picklauza and NCTC 12261. ΔCt was calculated by taking the average of each technical replicate. Genomic mean of all housekeeping genes (supplemental figure 2) was taken and then subtracted from the *qac* genes of interest. Average ΔCt between 3 biological replicates for NCTC 2261 was 0.58, and 0.73 for Picklauza.

### Identification of the *qacG* homolog in multiple oral streptococcal isolates highlights potential dissemination across IPF patient cohorts

PCR screening and sequencing of buccal *streptococcal* isolates recovered from additional patients with IPF clinical cohort identified the conserved *qacG* locus in three distinct isolates. All recovered sequences maintained complete conservation of the key residues maintained in the functional SMR *emrE* (Glu-14, Lys-22, Tyr-40, Trp-63, and His-110) establishing the *qacG* carriage is a mobile genomic feature rather than an isolated mutation (Figure 10).

**Figure 10.**
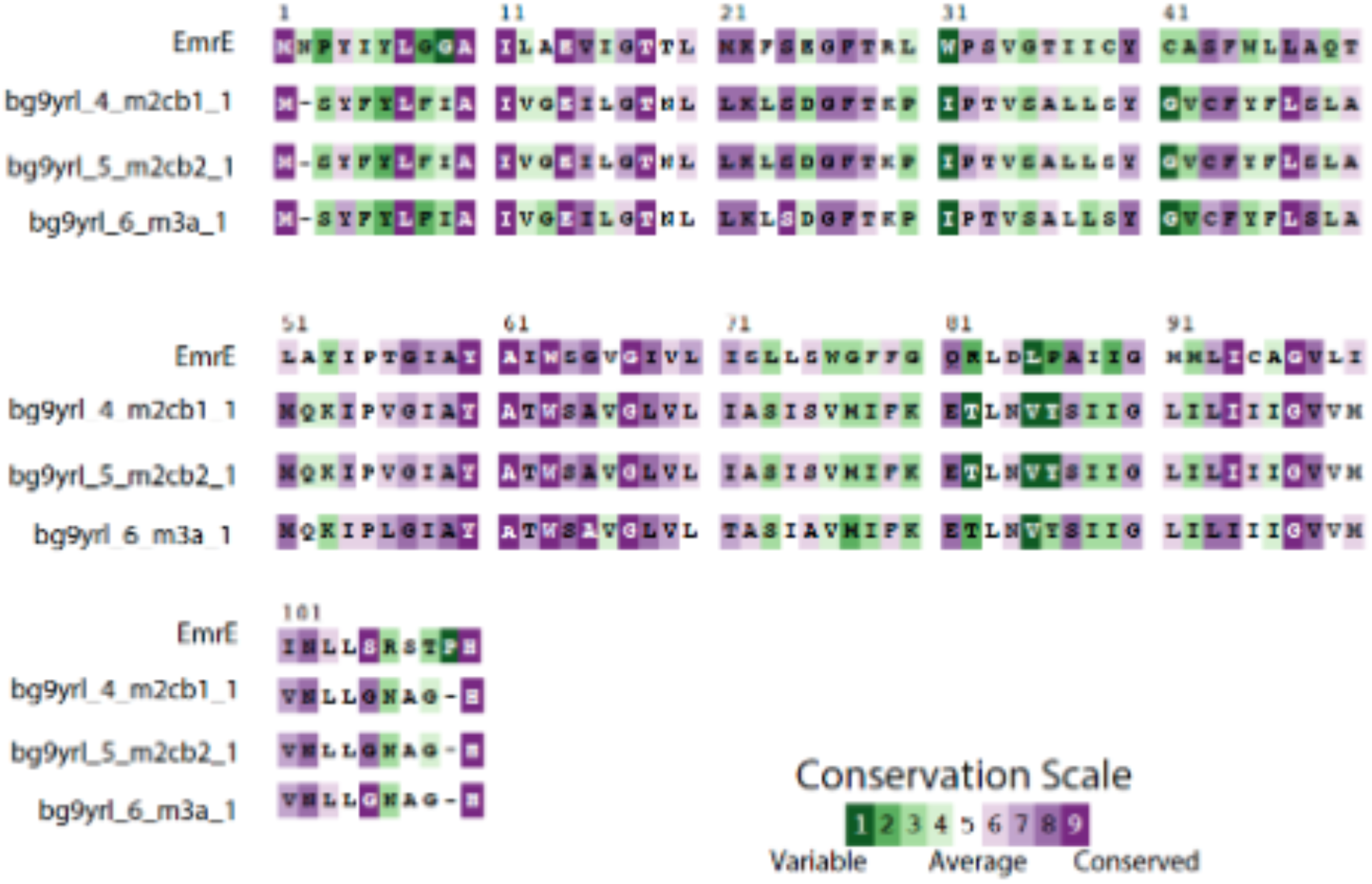
Multiple sequence alignment and conservation of *qacG* across *Streptococcus mitis* clinical isolates. Clustal-OMEGA amino acid sequence alignment comparing *qacG* genes from other *S. mitis* isolates from patients with IPF. compared against the E. coli EmrE multidrug transporter. The darkest shade of green (1) denotes highly variable amino acid positions, whereas dark purple (9) denotes most structurally and functionally conserved residues within the qacG sequence.

## DISCUSSION

We characterized Picklauza, a buccal *Streptococcus mitis* subsp. *mitis* isolate recovered from a patient with IPF and identified a *qacG* homolog predicted to encode an SMR-family efflux transporter. The locus was transcribed under standard growth conditions, retained residues conserved in characterized SMR proteins, and was located near genes associated with mobile genetic elements ^14,15,33^. Related *qacG* sequences were also detected in three additional oral isolates from patients with IPF. Despite these genomic and transcriptional features, Picklauza did not exhibit increased tolerance to BAC, BZT, or EtBr compared with NCTC 12261. The principal finding of this study is therefore a disconnect between the presence of a resistance-associated gene and the phenotype predicted from its annotation.

Several mechanisms could explain this genotype–phenotype discrepancy. First, basal transcript detection does not establish that QacG is translated at sufficient abundance or inserted correctly into the membrane. SMR transporters require appropriate topology, oligomerization, lipid interactions, and proton coupling, any of which may differ in the membrane environment of *S. mitis* ^34^. Second, the Picklauza protein may transport substrates that were not included in this study ^15,19^. Annotation as *qacG* is based primarily on sequence similarity, and substrate specificity cannot be inferred reliably from conserved residues alone. Third, transporter activity may depend on environmental conditions not reproduced in BHI broth, including pH, nutrient limitation, membrane stress, oxygen tension, or prior exposure to inducing compounds. Finally, increased efflux could be offset by differences in membrane permeability, energetic state, or other strain-specific susceptibility determinants ^13^.

The genomic context of *qacG* is consistent with possible horizontal acquisition. Genes annotated as an integrase homolog and a putative relaxase were located near the locus, and related *qacG* sequences were detected in additional isolates ^35^. Nevertheless, these observations do not themselves establish that the region is a complete, self-transmissible ICE or identify its donor lineage ^33,36^. Demonstrating horizontal transfer will require defining the element’s boundaries, comparing synteny across isolates and related streptococcal genomes, constructing gene and species phylogenies, and testing transfer experimentally.

Picklauza also differed from NCTC 12261 in aggregation, growth kinetics, carbon-source responses, and antimicrobial-response profiles. Picklauza formed visible aggregates in broth and consistently displayed capsular material by Anthony’s staining method ^30^. While NCTC 12261 also contains a capsule it was not always observed ^37^. Such traits could influence adherence, biofilm formation, or persistence on mucosal surfaces, although those functions were not measured directly. Picklauza grew more slowly than NCTC 12261 but reached a similar endpoint optical density and viable-cell yield, suggesting altered growth kinetics rather than reduced final carrying capacity under the tested conditions. Differences observed in Phenotype MicroArray assays further indicate physiological divergence between the strains, but targeted biochemical assays are needed to distinguish true substrate utilization from differential tolerance or assay-specific effects.

The recovery of Picklauza from a buccal swab is relevant to the possible contribution of oral communities to respiratory microbial exposure. Oral bacteria can enter the lower airways through microaspiration, and streptococci are commonly detected in both oral and respiratory samples ^5,34,38,39^. However, this study did not demonstrate that Picklauza colonized the lower respiratory tract or influenced IPF progression. The findings should therefore be interpreted as characterization of an oral isolate from a patient with IPF rather than direct evidence of adaptation to the fibrotic lung.

The QAC and EtBr results also require careful interpretation. BAC and BZT are used in a range of antiseptic, household, and personal-care formulations, whereas EtBr was included as a model SMR substrate rather than as a clinically relevant disinfectant exposure ^40^. Product-use concentrations cannot be compared directly with in vitro inhibitory concentrations because formulation, contact time, organic material, and delivery context substantially affect antimicrobial activity ^41–44^. Accordingly, the present data do not support conclusions about the effectiveness or safety of commercial products. Instead, they show that Picklauza did not gain measurable tolerance to the tested compounds merely by carrying and transcribing the annotated *qacG* locus.

The antimicrobial Phenotype MicroArray results likewise should be viewed as screening data. Notably, islands of resistance were present in the Picklauza profile that are absent in NCTC 12261. Lincosamide, Rifamycin, Amphenicol and Macrolides contain these putative islands of resistance ^45^. Picklauza showed distinct relative-growth patterns in several compound classes, but these assays did not use clinical breakpoints and do not establish clinical resistance. Variations in growth may reflect inducible responses, inoculum effects, technical variability, or concentration-specific physiology. Conventional broth microdilution with individual compounds and appropriate quality-control strains will be necessary to validate these patterns.

This study has several limitations that impact interpretation. Only a small number of candidate QacG substrates were tested, the concentration ranges were relatively coarse, and the laboratory growth conditions did not reproduce the oral or respiratory environment. Basal transcript detection was not accompanied by membrane-localization analysis, transport assays, gene deletion, or complementation. Further, the additional isolates were screened at the gene level, so their taxonomic identities, complete genomic contexts, and phenotypes remain to be established.

Future work should first validate the phenotype using standardized broth microdilution with narrower concentration intervals and additional SMR substrates, including cetyltrimethylammonium bromide and chlorhexidine. Comparing *qacG* expression before and after subinhibitory exposure could test inducibility. Protein abundance, membrane localization, intracellular accumulation of fluorescent substrates, and proton-coupling assays would further clarify why transcription does not produce the expected phenotype ^46^.

In summary, Picklauza carries and transcribes a *qacG* homolog with sequence and predicted structural features characteristic of SMR-family transporters, yet it remains sensitive to the tested QACs and EtBr. This genotype–phenotype mismatch cautions against inferring disinfectant resistance from gene annotation alone and establishes a framework for studying the regulation, substrate specificity, and mobility of SMR-family genes in oral *streptococci*.

## Supplementary Figures

**Supplementary Figure 1.**
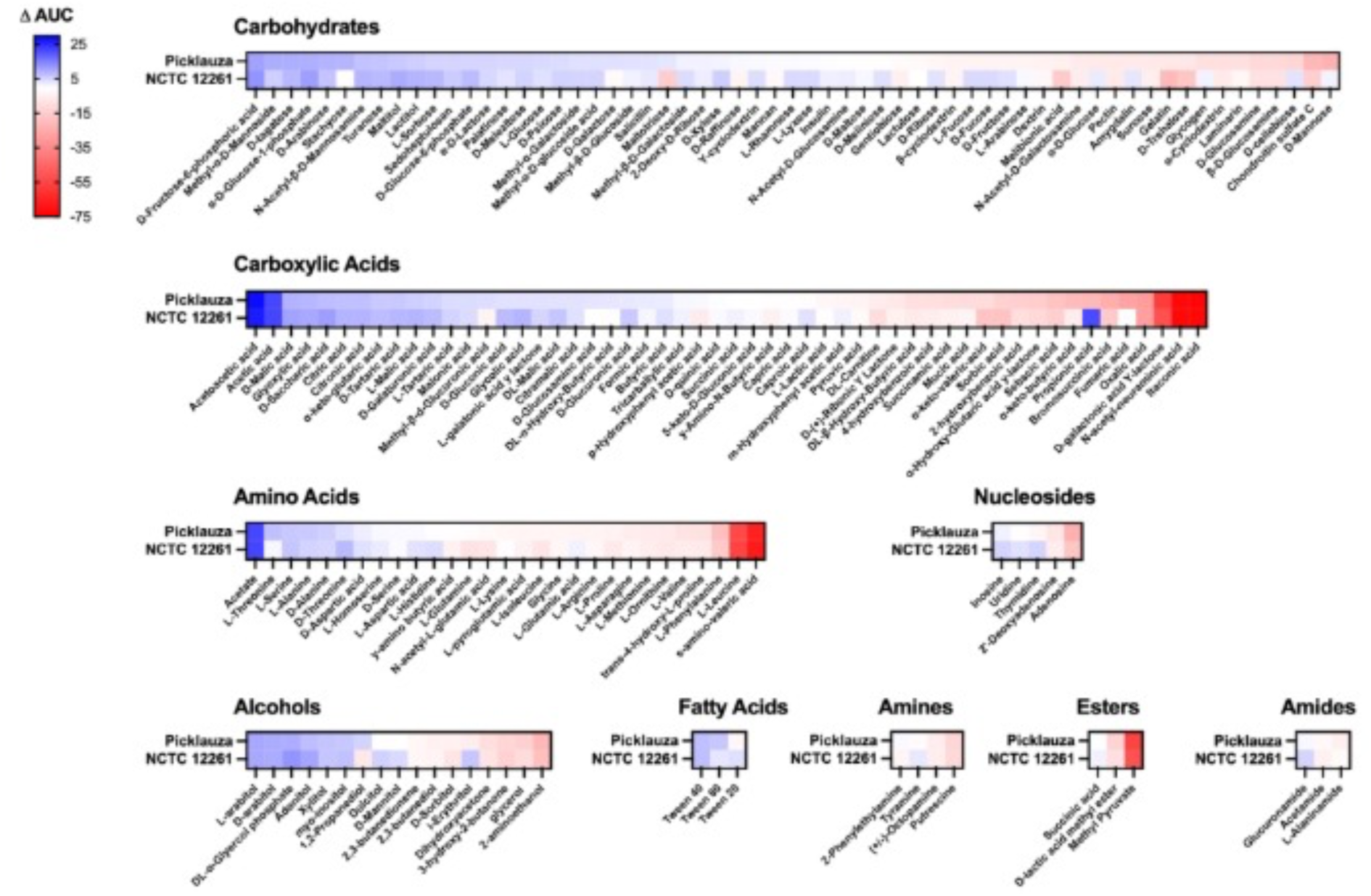
Phenotypic microarray profile of carbon source utilization in Picklauza and NCTC 12261. Heatmap containing carbon sources tested on NCTC 12261 and Picklauza. Ordered from highest ΔAUC (blue) to lowest (red) by Picklauza usage (n =3). Carbon profile determined via BIOLOG PM01 and PM02 plates. Cells inoculated in *Streptococcus* mixture IF and adjusted to OD 0.05 prior to incubation. AUC measured at OD740. ΔAUC calculated by subtracting the negative control from the average AUC value for each carbon source.

**Supplementary Figure 2.**
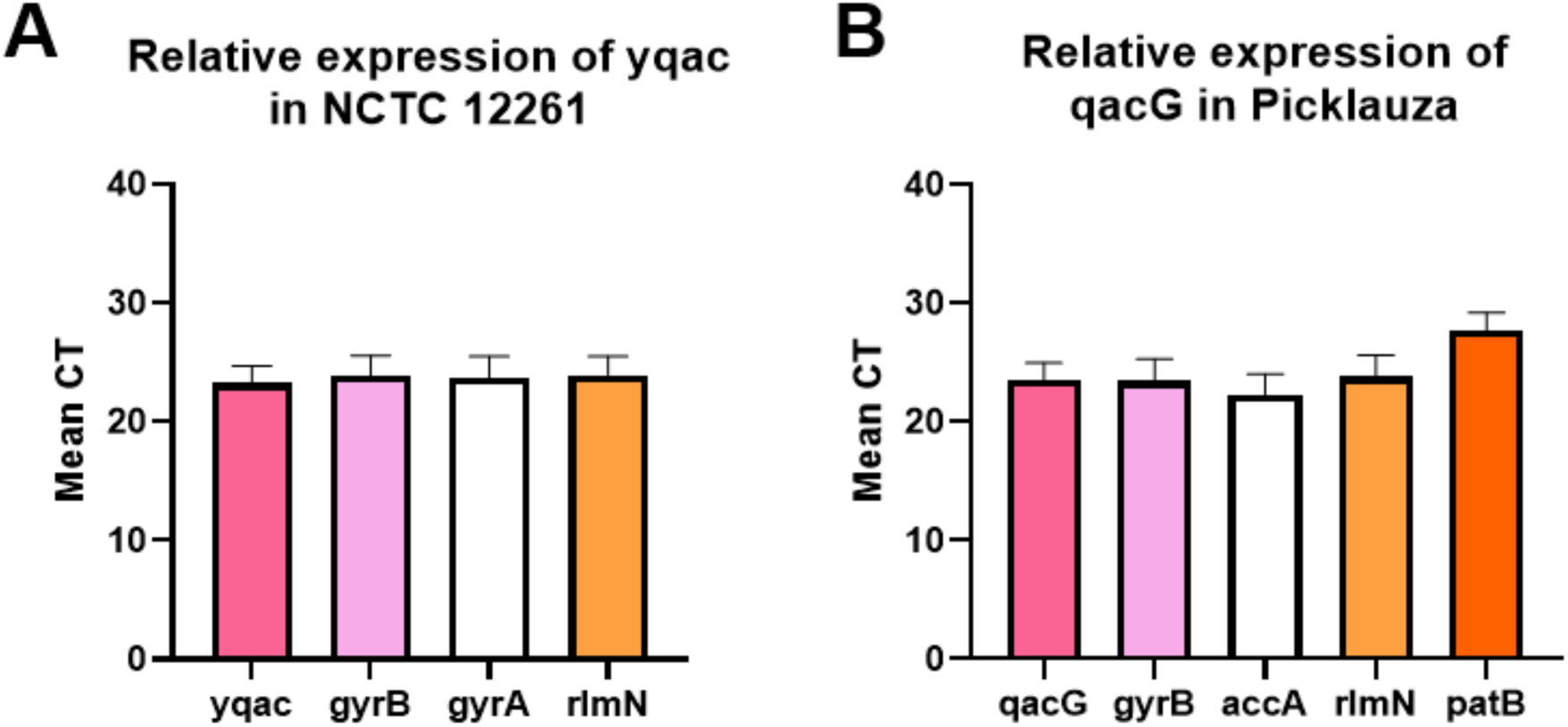
Mean CT values for Picklauza and NCTC 12261. A) Mean threshold cycle (CT) values of NCTC 12261 housekeeping genes and *yqac* target gene (dark pink). (B) Mean Ct values of Picklauza *qacG* target gene (pink). Data is represented as mean from 3 independent biological replicates.

**Supplementary Table 2.**
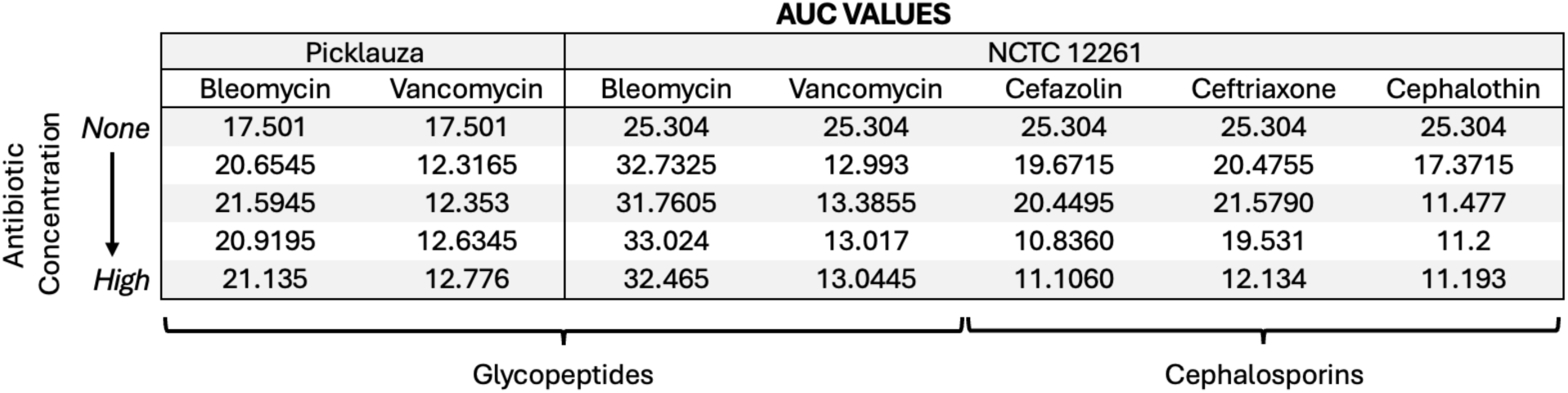
AUC specification for Picklauza and NCTC 12261 within specific antibiotic classes. Exact numerical AUC values (n=3) for select antibiotics included in Figure 4. Bleomycin and vancomycin (glycopeptide class) result in differential responses from both Picklauza and NCTC 12261 that are not clearly visible in Figure 4. Cefazolin, ceftriaxone, and cephalothin (cephalosporin class) show variable dose-dependent responses from NCTC 12261 that are not clearly visible in Figure 4.

|  |  | AUC VALUES |  |  |  |  |  |
| --- | --- | --- | --- | --- | --- | --- | --- |
| Antibiotic Concentration<br>None<br>↓<br>High | Picklauza |  | NCTC 12261 |  |  |  |  |
|  | Bleomycin | Vancomycin | Bleomycin | Vancomycin | Cefazolin | Ceftriaxone | Cephalothin |
|  | 17.501 | 17.501 | 25.304 | 25.304 | 25.304 | 25.304 | 25.304 |
|  | 20.6545 | 12.3165 | 32.7325 | 12.993 | 19.6715 | 20.4755 | 17.3715 |
|  | 21.5945 | 12.353 | 31.7605 | 13.3855 | 20.4495 | 21.5790 | 11.477 |
|  | 20.9195 | 12.6345 | 33.024 | 13.017 | 10.8360 | 19.531 | 11.2 |
|  | 21.135 | 12.776 | 32.465 | 13.0445 | 11.1060 | 12.134 | 11.193 |
|  |  |  | Glycopeptides |  | Cephalosporins |  |  |

